# The Effects of Physical Exercise on Osteocyte Lacunar Morphology and Orientation around Bioresorbable Implants

**DOI:** 10.64898/2026.10.29.762865

**Authors:** Irene Rodriguez-Fernandez, Thomas Bretschneider, Goran Lovric, Marco Stampanoni, Omer Suljevic, Nicole G. Sommer, Annelie-M. Weinberg, Uwe Y. Schwarze, Lukas Pircher, Christian Hellmich, Tilman A. Grünewald, Helga C. Lichtenegger, Andreas Menzel

## Abstract

In this study, we used synchrotron radiation micro-computed tomography (SR-µCT) to investigate the effect of early post-operative physical exercise at 2, 4, and 6 weeks on osteocyte lacunar distribution and morphology in a Sprague Dawley rat femoral defect model after implantation of a bioresorbable pin. Physical exercise has been shown to improve bone remodeling after fracture, but its effects on the lacuno-canalicular network in the presence of bioresorbable implants remain unclear. The structure of lacunae is linked to that of osteocytes, which play an important role in bone remodeling and adaptation to stress. We studied a large number of lacunae (around 670 000 per sample) in terms of their location, morphology, and orientation. We show that in newly formed bone, lacunae are larger and rounder with exercise, while in previously existing bone, lacunae are less oriented along the long bone axis when exercise is performed. In addition, close to the implant, lacunar density is higher with exercise in both newly formed bone and previously existing bone. These findings point to a higher degree of remodeling and a pronounced lacunar response to mechanical stimulation and may thus help clarify bone behavior with exercise around implants to better design post-surgical recovery treatments.

## 1 Introduction

Physical exercise plays a critical role in maintaining and enhancing bone health, offering significant benefits such as increased bone mass [1] and improved mechanical properties [2]. These effects are particularly vital during bone healing, where exercise supports the formation and adaptation of bone to restore its structural and functional integrity [3, 4]. Understanding these mechanisms is essential for optimizing rehabilitation strategies following fractures [5, 6].

Bone has a complex hierarchical structure over multiple length scales that is optimized for functionality and underlies its mechanical properties [7–11]. Bone contains several cell types, among which osteocytes are responsible for bone mechanotransduction [12–14] and play an important role in bone remodeling and adaptation to stress [15]. Osteocytes originate from osteoblasts trapped during bone formation and, thus, form a network embedded in the bone matrix. They reside in spaces called lacunae and connect to other osteocytes through thin channels named canaliculi. Studying lacunar distribution, shape, and orientation can reveal important indicators of bone quality and remodeling.

In the past, osteocytes were mainly studied with 2D techniques such as light microscopy [16–18] or electron microscopy [16, 19, 20]. However, due to the three-dimensional nature of the lacuno-canalicular network (LCN) and its inaccessibility within the mineralized bone matrix, three-dimensional measurements are essential. Details on 3D studies performed on lacunae can be found in multiple reviews comparing different imaging methods [21–24]. Although many of these methods, such as ptychographic X-ray computed tomography (PXCT) [25] and focused ion beam scanning electron microscopy (FIB/SEM) [24], provide very high resolution, they typically only allow the imaging of a few lacunae. To study the morphology and orientation of large lacunae populations, accurate techniques with a large field of view and micrometer resolution are required. Synchrotron micro-computed tomography (SR-µCT) allows for fast acquisition times and large fields of view, and it has been widely used in bone-lacunae studies [26–31]. Obata *et al*. [32] provide a comprehensive review of how to apply synchrotron microtomography to bone research.

After a bone fracture, orthopedic implants are often required to assist healing. While titanium is traditionally the material of choice, there is growing interest in bioresorbable implants, as their use can help avoid a second surgical intervention for implant removal [33]. In particular, Mg alloys have been widely studied as they minimize stress shielding, have excellent mechanical properties, and are biocompatible [34–36]. The lacuno-canalicular network in bone around bioresorbable Mg implants has been studied for some alloys, such as Mg-10Gd [37, 38], and PEO-coated Mg implants [39]. However, in these studies, the impact of physical stimuli was not investigated. It is, therefore, still an open question how physical exercise impacts the morphology of the lacuno-canalicular network around bioresorbable implants.

Here, we investigated the effect of physical exercise on the bone microstructure around bioresorbable implants with SR-µCT. Comparisons of exercised and non-exercised rat bone samples at 2, 4, and 6 weeks post-surgery revealed exercise-dependent differences in morphology and orientation of lacunae, indicative of increased remodeling activity.

## 2 Results

Each sample was divided into previously existing and newly formed bone (Figure 1). Previously existing bone corresponds to areas of bone present at the time of implantation whereas newly formed bone is bone that grew after. The regions were identified as described in the methods (section 5.4).

**Figure 1:**
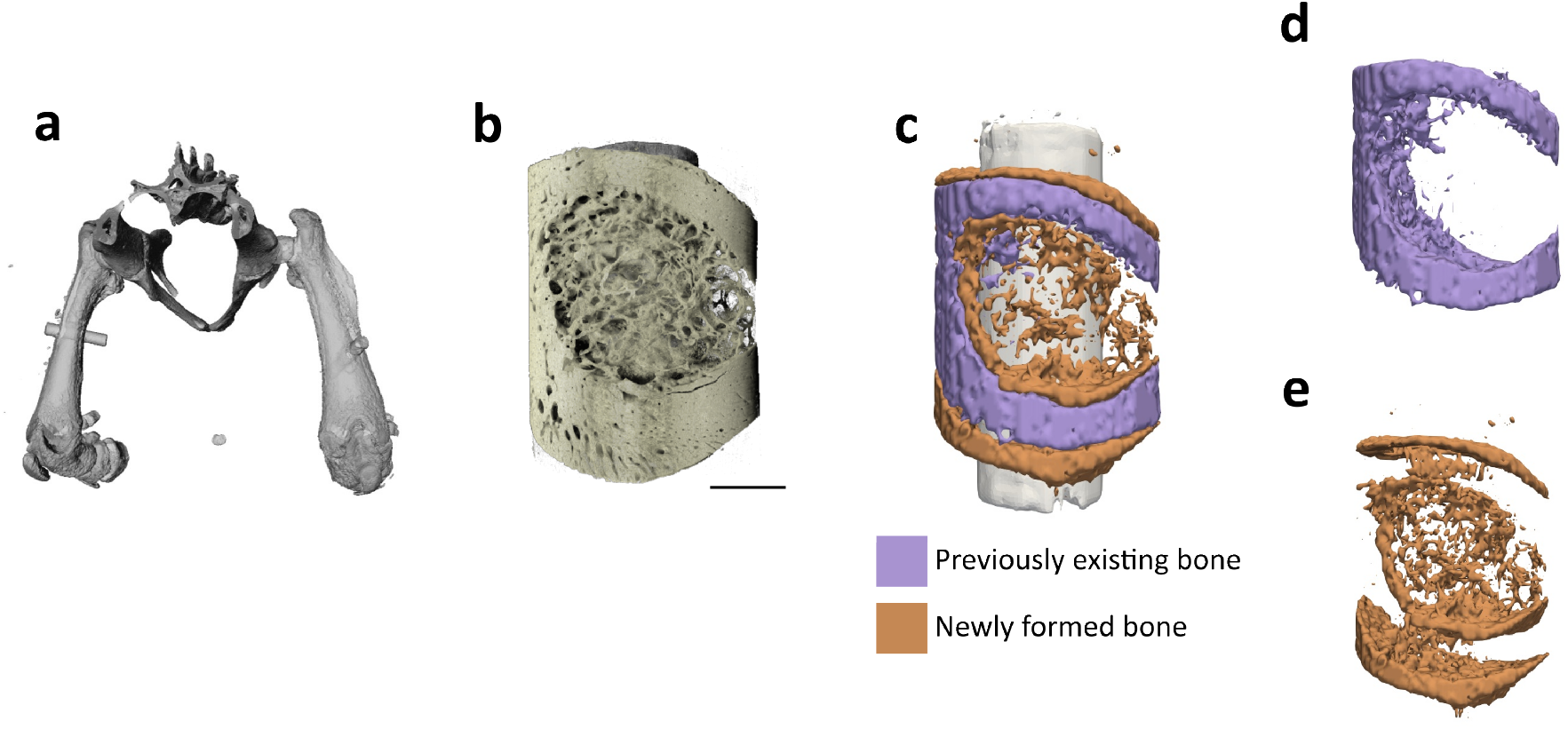
Experimental procedure. (a) *In vivo* µCT of the full hip with the implants. (b) SR-µCT reconstruction of 2w-Ex (scale bar of 1 mm). (c) Segmentation of previously existing and newly formed bone. (Adapted from [40]). (d) Previously existing bone mask. (e) Newly formed bone mask.

Of the six samples, the average number of lacunae per sample was 670 285, with a total number of analyzed lacunae of 4 021 710, 50–75 % of which were found in previously existing bone, depending on the sample. For details per sample, see Tables S1 and S2.

### 2.1 3D morphology of lacunae

Representative slices of the SR-µCT reconstructions are shown in Figure 2. Differences in lacunar morphology between previously existing and newly formed bone are evident in Figure 2b. Furthermore, in previously existing bone, noticeable changes in lacunar morphology and orientation were observed close to the implant, within the first 200 µm (Figure 2c–d). This distance-dependent behavior is also reflected in the quantitative analysis of lacunar parameters as a function of distance from the implant (see Figures S1 and S2). The influence of the implant appeared to be confined to approximately 100–200 µm from the implant interface for both previously existing and newly formed bone, beyond which the curves are largely flat. For this reason, in previously existing bone, lacunae were further subdivided into regions up to 200 µm from the implant interface and regions farther away. This distinction was not applied to newly formed bone, as all tissue of this type developed under the influence of the implant. In addition, previously existing bone at 6 weeks located farther than 200 µm from the implant was used as a reference, as it represents the most mature and least altered bone in this study.

**Figure 2:**
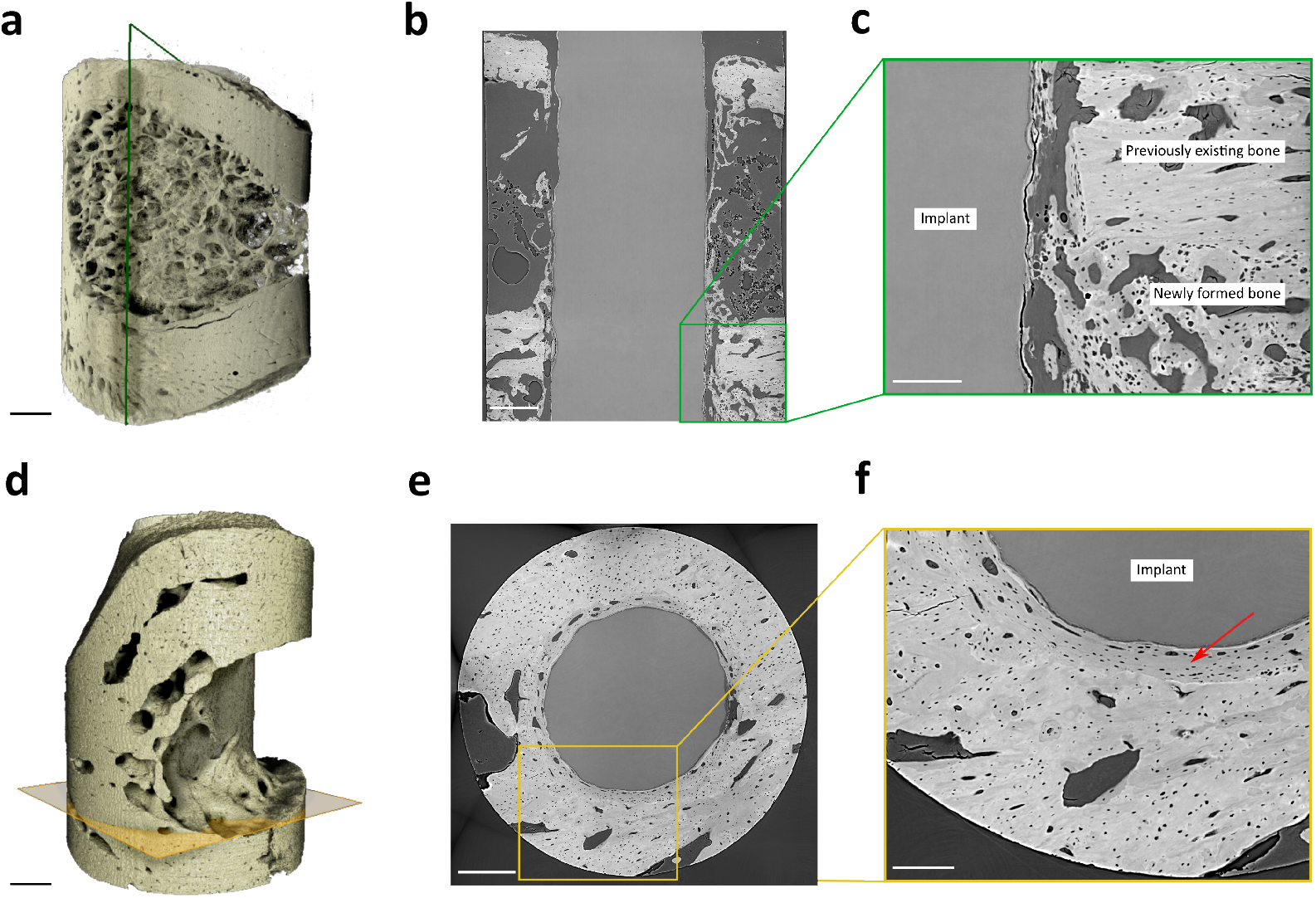
Reconstruction slices. (a) SR-µCT reconstruction of 2w-Ex. (b) Reconstructed slice along the implant shaft and long bone axis (green plane in a). (c) Zoomed-in image of at the interface between previously existing and newly formed bone regions. (d) SR-µCT reconstruction of 6w-Ex. (e) Reconstructed slice perpendicular to the implant shaft (orange plane in d), corresponding to previously existing bone. (f) Zoomed-in image of (e). Scale bars: 500 µm (a,b,d,e), 200 µm (c,f).

Grouping lacunae according to bone type and distance from the implant and evaluating the morphological parameters for each individual sample allowed us to follow changes in lacunar morphology with bone type and position. Figure 3 shows the distributions of lacunar morphological parameters in newly formed bone. Non-exercised samples (nEx) are shown in blue and exercised samples (Ex) in orange; dashed lines indicate median values (see also Tables S1 and S2).

**Figure 3:**
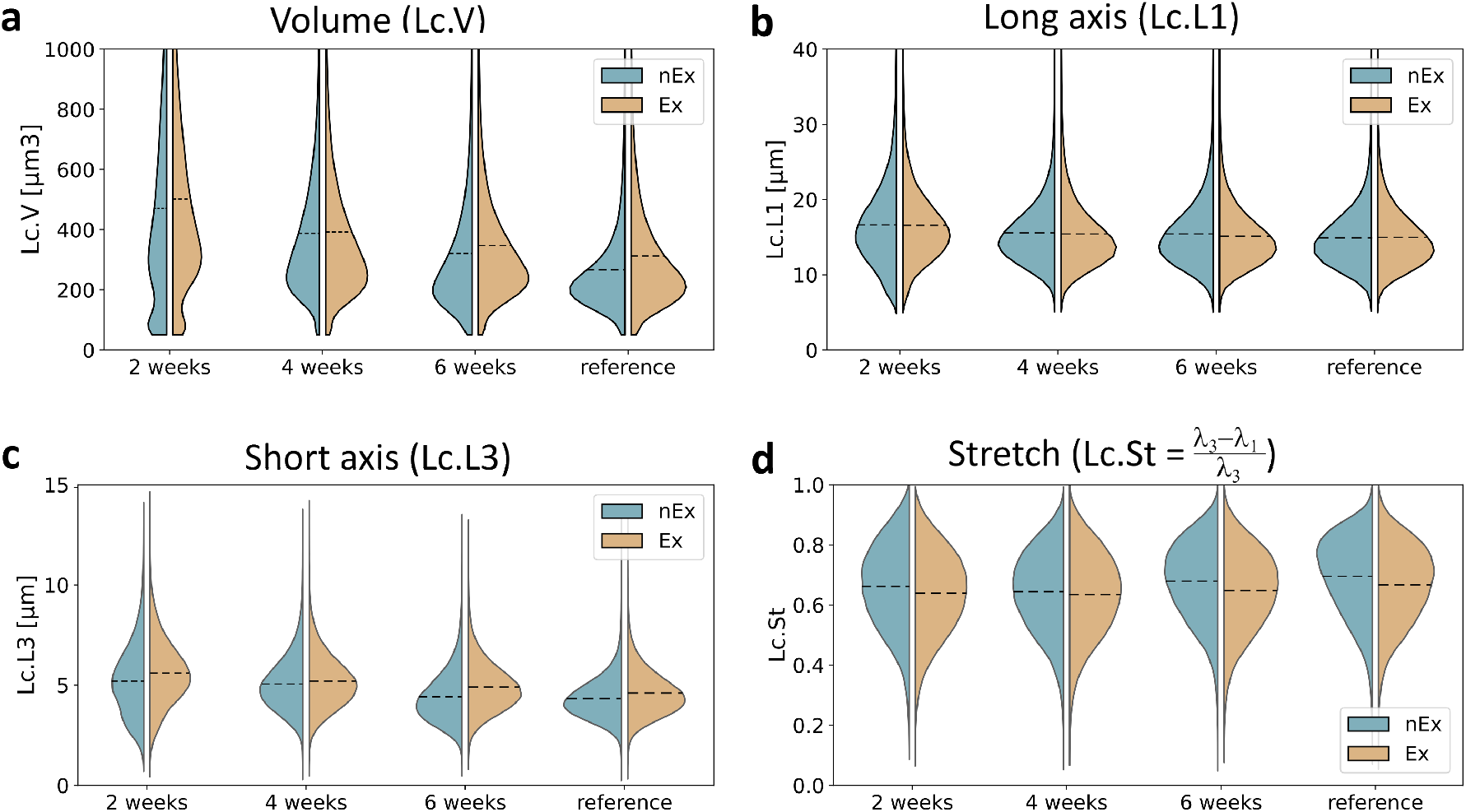
Distributions of lacunar morphological properties in newly formed bone. Plots comparing the distributions of Lc.V (lacunar volume) (a), Lc.L1 (long axis) (b), Lc.L3 (short axis) (c), and stretch (d), for non-exercised (nEx, blue) and exercised (Ex, orange) samples. The dashed line represents the median value of the distribution. The reference corresponds to previously existing bone at 6 weeks, farther than 200 µm from the implant.

In newly formed bone, the median lacunar volume (Lc.V) at 2 weeks was 387 µm^3^ in nEx and 422 µm^3^ in Ex (Figure 3a). With time, volumes decreased to 260 µm^3^ (nEx) and 296 µm^3^ (Ex) at 6 weeks. At all time points, volumes were larger in exercised samples. Moreover, the volume distributions were broader at 2 weeks and progressively narrowed over time, approaching the reference values.

The lacunar long axis (Lc.L1) exhibited a similar trend (Figure 3b). At 2 weeks, Lc.L1 measured 15.92 µm in both nEx and Ex samples and decreased with time. At 6 weeks, Lc.L1 was 14.74 µm in nEx and 14.45 µm in Ex. The short axis (Lc.L3) also decreased with time, from 5.11 to 4.26 µm in nEx and from 5.52 to 4.80 µm in Ex (Figure 3c). Across all time points, Lc.L3 was larger in Ex samples.

Consistent with these axis changes, stretch values were lower in Ex samples (Figure 3d), although the temporal trend was less pronounced. Together, the combined trends in volume, axis lengths, and stretch indicate that lacunae in newly formed bone are less elongated (i.e., geometrically rounder) under exercise and that their distributions become more homogeneous over time. For previously existing bone, the corresponding distributions are shown in Figure S3. In this compartment, no systematic trends with exercise or time were observed.

### 2.2 Lacunar density

Lacunar density (Lc.D) has been reported as an indicator of remodeling activity [41]. The variation of Lc.D with time and exercise is shown in Figure 4. In newly formed bone (Figure 4a), the lacunar density was higher in exercised samples, with the largest difference at 2 weeks (60 233 mm^*−*3^ (nEx) and 67 998 mm^*−*3^ (Ex)). At later time points, the lacunar density increased relative to 2 weeks, with only small differences between exercise and non-exercise.

**Figure 4:**
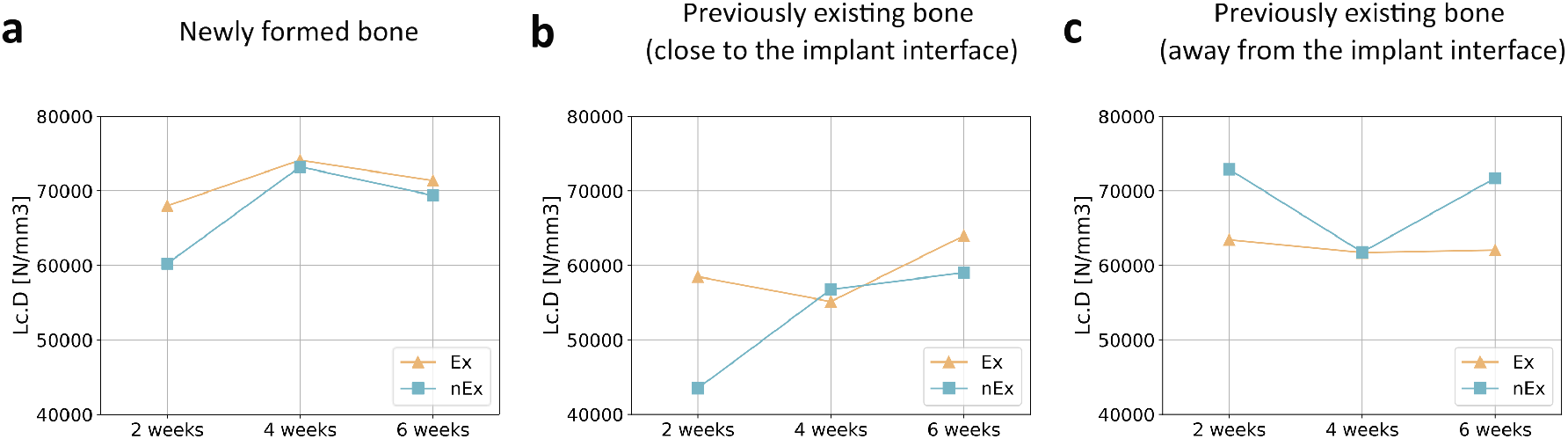
Lacunar density trends with time and exercise. (a) Newly formed bone. (b) Previously existing bone, up to 200 µm to the bone–implant interface. (c) Previously existing bone, farther than 200 µm. Non-exercised (nEx) samples are presented in blue and exercised (Ex) in orange.

As before, for previously existing bone, the lacunae were further subdivided by distance from the implant into bone up to 200 µm from the implant interface and bone farther than that. Close to the implant interface (Figure 4b), the lacunar density was generally higher with exercise. However, far from the implant (Figure 4c), the lacunar density was lower in the exercised samples, with values remaining approximately 62 000 mm^*−*3^ over time.

### 2.3 Lacunar orientation

Osteocytes, and therefore lacunae, are expected to be oriented along the long bone axis [42]. Here, we quantified lacunar orientation by computing the angle between the long bone axis and the eigenvector corresponding to the longest lacunar axis. This angle was obtained from the dot product of the two unit vectors representing these directions. Results are shown in Figure 5 (previously existing bone) and Figure 6 (newly formed bone).

**Figure 5:**
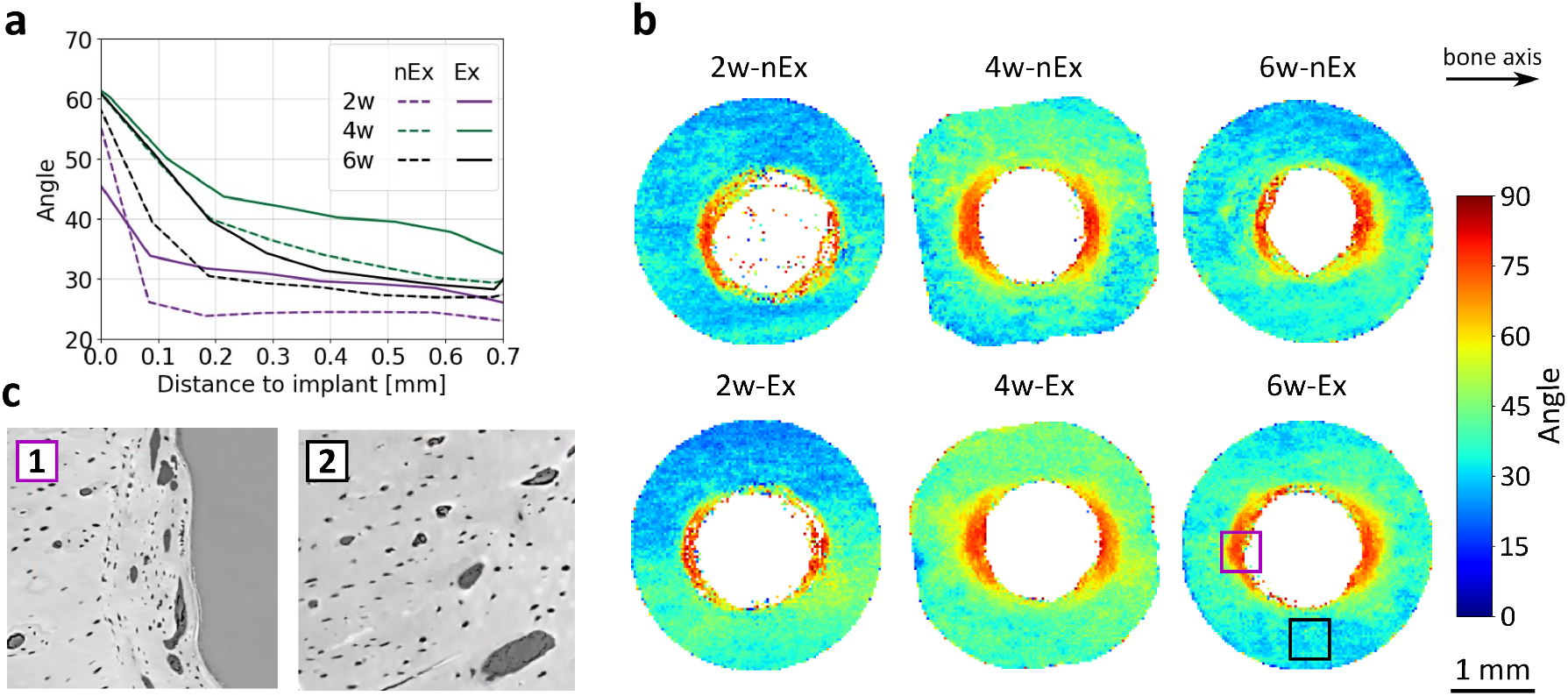
Lacunar orientation in previously existing bone. (a) Angle between the longitudinal bone axis and the lacunar orientation as a function of distance from the bone–implant interface. (b) Median of the lacunar orientation along the implant direction for non-exercised (nEx, top row) and exercised (Ex, bottom row) samples. The color represents the same angle as in (a). Reconstructed slices at the bone–implant interface (1) and far from the implant (2).

**Figure 6:**
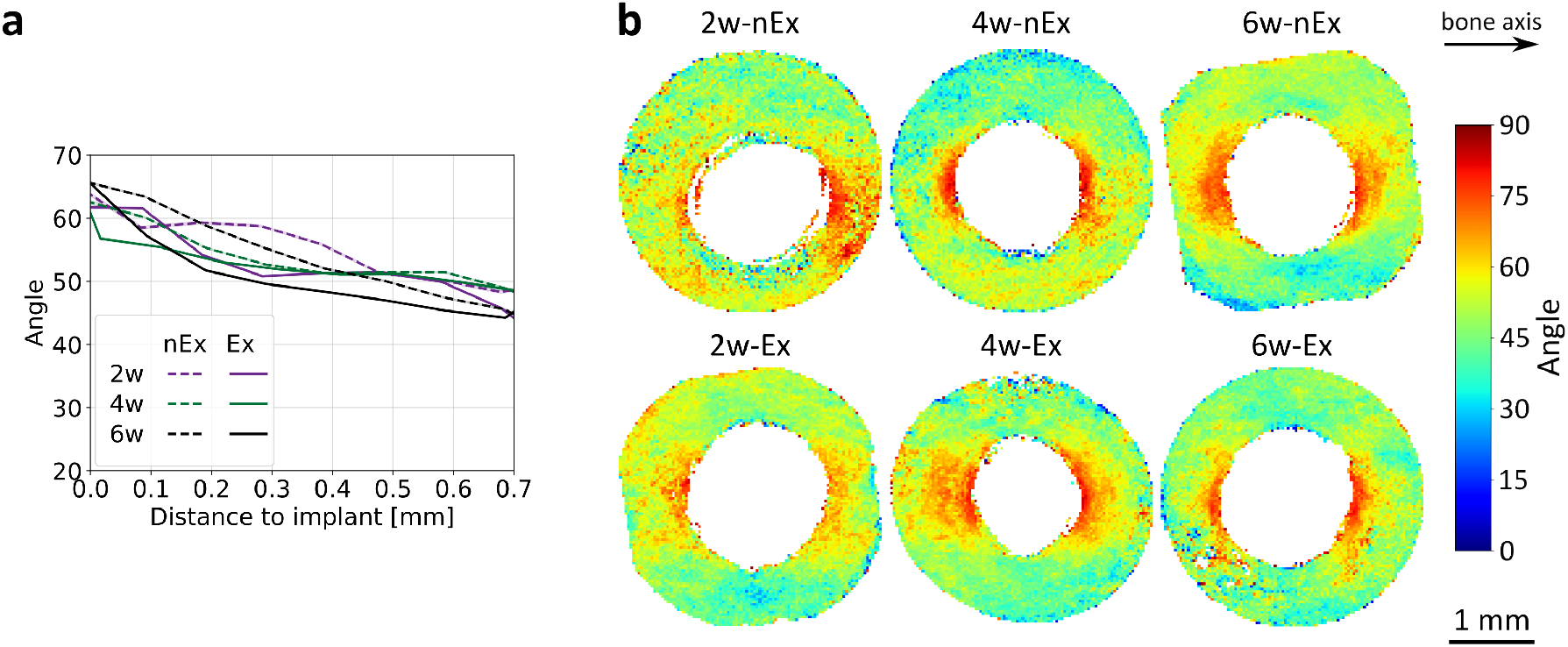
Lacunar orientation in newly formed bone. (a) Angle between the longitudinal bone axis and the lacunar orientation as a function of distance from the bone–implant interface. (b) Median of the lacunar orientation along the implant direction for non-exercised (nEx, top row) and exercised (Ex, bottom row) samples. The color represents the same angle as in (a).

In Figure 5a, this angle is plotted as a function of distance from the bone–implant interface. Lacunar orientation showed a large deviation from the long bone axis at the interface, and a gradual reduction over the first 100–200 µm from the implant for all samples. Farther from the implant, lacunae were predominantly oriented along the long bone axis. The reduction in angle within the first 200 µm was more pronounced in nEx than in Ex samples, and overall deviations from the bone axis were larger in Ex than in nEx samples at the same time point.

This behavior was also evident in Figure 5b, which visualizes the spatial distribution of the median orientation angle (computed per voxel from the lacunae contained in that voxel) along the implant direction. Dark blue indicates lacunae aligned with the long bone axis, whereas red indicates orientations close to 90^*°*^. In all samples, regions close to the implant contain areas with angles approaching 90^*°*^ (red), while farther from the implant the angles were typically in the range 20–40^*°*^. Figure 5c illustrates the reorientation of lacunae following the implant surface close to the interface (1) and alignment along the long bone axis farther from the implant (2).

In newly formed bone (Figure 6a), the angle decreased with distance from the implant, and the overall deviation from the bone axis was larger than in previously existing bone. Larger regions close to the implant showed reorientation following the implant surface (Figure 6b). No pronounced effect of exercise was apparent in this compartment.

## 3 Discussion

The present study was designed to determine the effect of physical exercise on the osteocyte lacunar network around Mg-based bioresorbable implants. Using SR-µCT, we imaged six large bone volumes (3 mm *×* 3 mm *×* 7 mm) and analyzed approximately 670 000 lacunae per sample, representing, to our knowledge, the largest lacunar population studied with SR-µCT to date. We identified differences in lacunar morphology and orientation between exercised and non-exercised samples in both previously existing and newly formed bone.

### 3.1 Influence of exercise on the morphology

Osteocytes play an important role in mechanotransduction and bone remodeling [14], and the shape, orientation, and density of lacunae affect osteocyte function [15]. Lacunar density depends on bone type, age, and loading status [37]. In this work, the average lacunar density was approximately 69 000 mm^*−*3^ in newly formed bone and 63 000 mm^*−*3^ in previously existing bone. These values agree well with previous lacunar studies [26, 27, 43]. We reported higher lacunar density with exercise in newly formed bone and in previously existing bone close to the implant (Figure 4a,b). Higher density has been suggested to indicate regions of increased remodeling activity [41].

Previous studies reported mean lacunar volumes of 200–500 µm^3^ [27, 37, 44] and dimensions of approximately 3 µm *×* 9 µm *×* 15 µm [21, 26]. In this study, we obtained a mean lacunar volume per sample of 300–500 µm^3^, and lacunae were fitted by an ellipsoid with dimensions of approximately 5 µm *×* 8 µm *×* 15 µm, in agreement with previously reported values.

In newly formed bone, we observed larger and rounder lacunae compared to previously existing bone. This bone forms in response to the fracture, where osteoblasts rapidly deposit collagen fibers. As the matrix develops, the cells become entrapped and differentiate into osteocytes [45]. Initially, healing bone corresponds to woven bone, where lacunae are rounder and larger [24]. With time, this type of bone develops into lamellar bone. Accordingly, lacunae became smaller and more stretched with time, approaching the reference values of previously existing bone at 6 weeks, far from the implant (Figure 3). Lacunae in newly formed bone were also larger and rounder with exercise at all time points. Osteocyte shape impacts cellular elasticity, and round osteocytes have been shown to be less stiff and respond more to mechanical stimulation [46].

In previously existing bone, which was already lamellar at the time of implantation and experiences minimal remodeling, lacunae were smaller and more stretched. Physical exercise did not significantly affect lacunar shape in this compartment. The differences in lacunar morphology between both bone types are illustrated in Figure 2b.

### 3.2 Influence of exercise on the orientation

Microstructural and nanostructural orientations in bone are correlated. The orientation of the main lacunar axis and that of the surrounding collagen fibers have been shown to coincide [42, 47]. This orientation is influenced by the loading state; in long bones such as the femur, collagen fibers tend to align along the main bone axis [27, 48].

In our previous study on how physical exercise affects the nanostructure of bone around bioresorbable implants [40], we analyzed the orientation of mineralized collagen fibers in the same samples. In that study, it was assumed that the mineral is co-aligned with the fibrillar direction of the collagen, as demonstrated previously [49]. In agreement with previous studies [50–52], we observed there that, in previously existing bone, the mineralized collagen fibers wrapped around the implant close to the bone–implant interface and aligned predominantly along the main bone axis farther away. In the present study, lacunae in previously existing bone were likewise predominantly oriented along the long bone axis far from the implant. Close to the implant, however, reorientation following the implant surface was observed (Figure 5b). This reorientation is consistent with the altered mechanical environment induced by implantation, with high shear stresses occurring close to the implant surface, which are likely to induce significant bone remodeling and, therefore, deviate the lacunae from the main bone axis [53]. Moreover, lacunae were overall more aligned with the bone axis in non-exercised samples (Figure 5a). These findings agree well with the fiber alignment we observed previously.

In woven (newly formed) bone, collagen fibers are deposited rapidly and with little preferential orientation. In our previous study [40], we also observed that the mineralized collagen fiber orientation was more random at early time points, and that wrapping around the implant was more pronounced and extended over larger regions than in previously existing bone. Similarly, lacunae in newly formed bone exhibited long-axis orientations consistent with a wrapping pattern around the implant and showed weaker alignment with the long bone axis far from the implant (Figure 6b). However, the effect of exercise on orientation was less pronounced in newly formed bone (Figure 6a).

All in all, our results suggest that physical exercise has a significant impact on the morphology and orientation of the lacuno-canalicular network and, therefore, on the bone mechanotransduction and remodeling around bioresorbable implants.

We acknowledge several limitations of this study. In particular, only one specimen per group could be investigated with SR-µCT. This is due to the long measurement times to scan large volumes at high resolution and the limited access to synchrotron experiments. Nevertheless, each sample encompassed the analysis of very large lacunar populations (490 000–850 000 lacunae), ensuring statistical robustness. A further limitation is that only one exercise protocol was investigated; other protocols may yield different outcomes. Similarly, only XHP-Mg implants were examined, and responses to other implant materials in the presence of exercise remain to be explored.

## 4 Conclusion

In this work, we segmented and quantified a large lacunar population (on average approximately 670 000 lacunae per sample) using SR-µCT and established an analysis pipeline capable of processing such data volumes reliably and efficiently. We found that physical exercise is associated with changes in lacunar density, morphology, and orientation in both previously existing and newly formed bone. In newly formed bone, exercise resulted in larger and rounder lacunae, suggesting an enhanced response to mechanical stimulation. Lacunar density was higher with exercise in newly formed bone and in previously existing bone close to the implant, consistent with increased remodeling activity. In previously existing bone, lacunae were less aligned with the long bone axis under exercise, in agreement with the nanostructural mineralized collagen orientation observed in our previous study.

These findings provide insight into the microstructural response of bone to mechanical stimulation in the presence of bioresorbable implants and may help inform post-surgical rehabilitation strategies and assessments of mechanical adaptation. The analytical framework developed here further enables future high-throughput investigations of lacunar organization in large bone volumes.

## 5 Materials and Methods

### 5.1 Samples

All animal experiments were approved by the Austrian Federal Ministry for Science and Research under permit BMWFW-2020-0.363.120 and were carried out in accordance with the European Convention for the Protection of Vertebrate Animals Used for Experimental and Other Scientific Purposes. Male Sprague Dawley rats underwent surgery at 6 weeks of age. The surgery protocol is described in detail in Suljevic *et al*. [4]. In brief, the surgical area was cleaned with alcohol, and transverse skin incisions were made on the femoral diaphyses. A surgical drill with a 1.55 mm bur was then used to create a cylindrical hole at the diaphysis, and XHP-Mg [54] pins were implanted. Absorbable sutures were used to close the skin. All rats received subcutaneous injections of Carprofen (200 mg*/*kg; Rimadyl, Pfizer Corporation, Vienna, Austria) for analgesia. After surgery, *in vivo* µCT scans were performed.

The rats were randomly divided into two groups, those assigned to undergo treadmill exercise (Ex) and those allowed to move in an unrestrained way (nEx). Their behavior was monitored daily until the experiment concluded. Two days after the implantation, Ex rats underwent a 2-week (5 days per week) treadmill running exercise, an exercise protocol chosen because of its positive effects on bone regeneration and osteogenesis [4]. The animals were then euthanized at 2, 4, and 6 weeks after implantation.

After explantation, the bone was fixed, dehydrated, and embedded in PMMA (Technovit 9100 New, Heraeus Kulzer, Wehrheim, Germany). The samples were then machined into cylindrical shapes using a precision saw (Struers Accutom 50, Struers, Ballerup, Denmark). The samples analysed in this study were the same specimens previously investigated using SAXS tensor tomography (SASTT) in [40].

### 5.2 *In vivo* micro-computed tomography (µCT)

*In vivo* micro-computed tomography of femoral bones was performed using a Bruker Skyscan (Bruker Skyscan, Kontich, Belgium) directly after surgery. The animals were positioned in a supine position during the scan with the femurs oriented at a 90^*°*^ angle. The image reconstruction was performed using NRecon software (Skyscan, Kontich, Belgium). The *in vivo* µCT data was used to discriminate previously existing bone and bone formed after the implantation (Section 5.4).

### 5.3 Synchrotron phase-contrast computed tomography (SR-µCT)

SR-µCT experiments were performed at the X02DA TOMCAT beamline of the Swiss Light Source SLS at the Paul Scherrer Institute PSI, Villigen, Switzerland [55]. The X-ray beam at TOMCAT was produced with a 2.9 T superbending magnet from a 2.4 GeV storage ring with 400 mA ring current, operated in top-up mode. The measurements were conducted at an X-ray beam energy of 21 keV, selected with a double-multilayer monochromator (DMM). A scientific CMOS detector with 6.5 µm pixel size (pco.Edge) was coupled to a 20 µm thick LuAG:Ce scintillator and a 20 *×* magnifying visible-light optics, resulting in an isotropic voxel size of 0.325 µm. The sample was placed 25 m from the X-ray source, and the sample-to-detector distance was set to 38 mm, yielding moderate edge enhancement while remaining in the Fresnel near-field regime to allow for single-distance phase retrieval. After flat-field correction, Paganin phase retrieval [56] was performed on the single tomographic projections, followed by *gridrec*-based CT reconstructions [57]. For each scan, 3000 projections over 360^*°*^ were acquired with a 170 ms exposure time. Since the samples were larger than the available field-of-view (FOV), a total of 32–72 subvolumes were scanned per sample. The subvolumes were selected to have approximately 30 % overlap and all volumes of a given scan were then input to a non-rigid stitching algorithm to produce high-resolution and high-FOV volumes [58].

### 5.4 Segmentation into bone types

To distinguish between bone present at the time of implantation and bone formed subsequently, the reconstructed volumes were partitioned into two bone compartments: previously existing bone and newly formed bone. Previously existing bone corresponds to regions already present at the time of surgery, whereas newly formed bone represents tissue that developed after implantation.

Operationally, these compartments were identified by registering the *ex vivo* SR-µCT data to the *in vivo* µCT scans acquired immediately after surgery. For this purpose, the SR-µCT was binned to match the voxel size of the *in vivo* µCT (40 µm), and the two datasets were overlaid using Paraview software [59], guided by matching morphological features. Voxels containing bone signal in both datasets were classified as previously existing bone. In contrast, voxels showing bone only in the SR-µCT data, but not in the *in vivo* µCT, were classified as newly formed bone, reflecting bone growth after implantation [40].

### 5.5 Image post-processing – Segmentation of lacunae

Due to the large amount of data to process, lacunae were segmented on binned datasets, reducing the voxel size by a factor of two to 0.65 µm, and dividing each dataset into 50 subvolumes to enable processing of smaller data volumes. The segmentation workflow is illustrated in Figure 7 and explained as follows. First, a volume-of-interest (VOI) is selected from the CT-reconstructed and stitched volume data (Figure 7a), after which low and high thresholds were applied to differentiate bone from implant and voids, including blood vessels, lacunae, etc. (Figure 7d). Each void was then labeled using ndimage.label from SciPy, which assigns a unique label to connected components in a binary image (Figure 7e). Voids intersecting the subvolume boundaries were removed to exclude incomplete lacunae from further evaluation. Subsequently, a size threshold was applied, and only voids with volumes of 50–2000 µm^3^ were classified as lacunae [21] (Figure 7f). A three-dimensional rendering of the segmented lacunae is shown in Figure 7g. Finally, segmented lacunae were assigned to either of the previously or newly formed bone compartments, based on their spatial location with the corresponding bone masks defined above.

**Figure 7:**
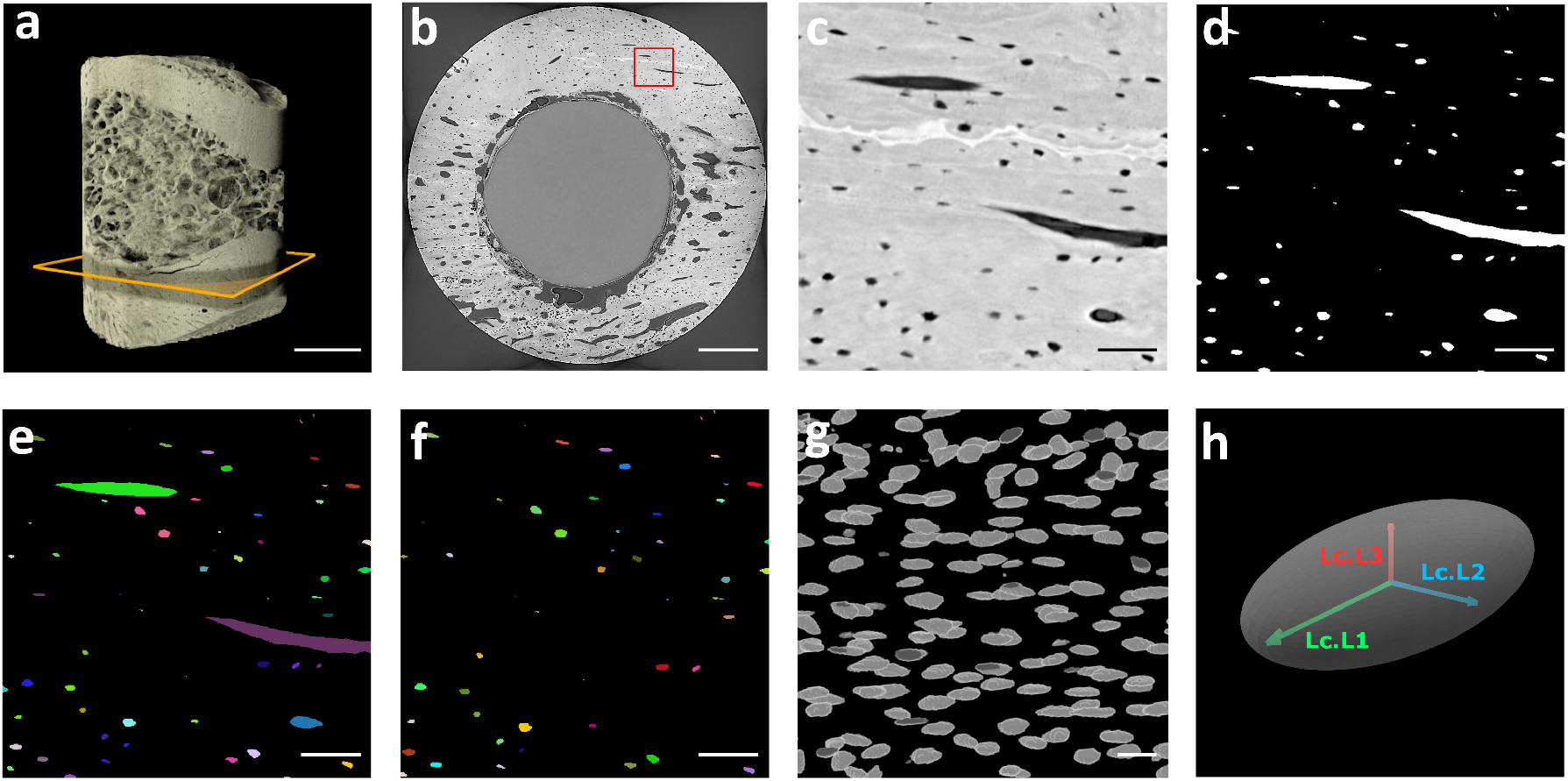
Image post-processing and segmentation. (a) Binned sample reconstruction. (b) Reconstructed slice, corresponding to the yellow square in (a). (c) Zoomed-in view of the slice, corresponding to the red square in (b). (d) Binary segmented slice. (e) Labeled slice. (f) Labeled lacunae. (g) 3D rendering of the segmented lacunae. (h) Fitting of an ellipsoid to the lacunae, with the axes Lc.L1 (long), Lc.L2 (medium), and Lc.L3 (short). Scale bars: 1 mm (a), 500 µm (b), 50 µm (c,d,e,f), 15 µm (g).

Bone volume (BV) was defined as the product of the voxel volume and the number of voxels classified as bone within the previously defined bone compartments. For this calculation, lacunae were included in the bone mask, such that only voids larger than 2000 µm^3^, corresponding to larger structures such as the implant, and blood vessels, were excluded.

### 5.6 3D lacunar descriptors

Morphological and orientation descriptors were extracted for each lacuna to characterize their geometry and spatial organization. Descriptor extraction was performed using the Python-based open-source software SPAM (Software for Practical Analysis of Materials) [60]. For each lacuna, the bounding box, center of mass, and moments of inertia were computed. Lacunar orientation was determined from the eigenvalues (*λ*_3_, *λ*_2_, *λ*_1_) and eigenvectors of the inertia tensor, with *λ*_3_ *≥ λ*_2_ *≥ λ*_1_. Lacunar shape was characterized by the stretch 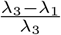, as introduced by Mader *et al*. [61]. Lacunar volume (Lc.V) was calculated by counting the voxels associated with each lacuna.

Assuming an ellipsoidal lacunar shape, the lacunar dimensions (Lc.L1, Lc.L2, Lc.L3) were estimated from the principal moments of inertia following the approach described by Ikeda *et al*. [62], where Lc.L1 corresponds to the longest axis, Lc.L2 to the intermediate axis, and Lc.L3 to the shortest axis. A schematic of the fitted ellipsoid and the corresponding descriptors is shown in Figure 7h.

Lacunar density (Lc.D) was defined as the total number of lacunae in a sample divided by its bone volume (Lc.N/BV).

Table 1 summarizes the evaluated parameters.

**Table 1:**
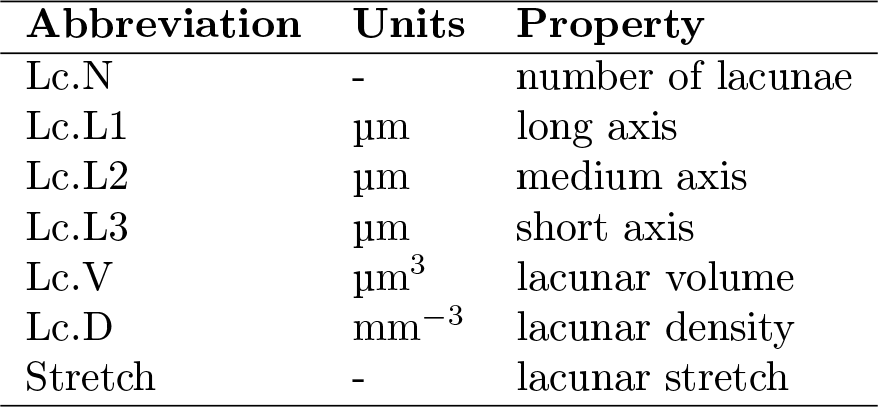
Lacunar descriptors.

| Abbreviation | Units | Property |
| --- | --- | --- |
| Lc.N | - | number of lacunae |
| Lc.L1 | $\mu\text{m}$ | long axis |
| Lc.L2 | $\mu\text{m}$ | medium axis |
| Lc.L3 | $\mu\text{m}$ | short axis |
| Lc.V | $\mu\text{m}^3$ | lacunar volume |
| Lc.D | $\text{mm}^{-3}$ | lacunar density |
| Stretch | - | lacunar stretch |

### 5.7 Distance from implant analysis

During the SR-µCT measurements, the implant axis was aligned with the Y-axis of the experimental setup. From the SR-µCT reconstruction, virtual cross-sectional slices in the X-Z plane were extracted at both ends of the implant (e.g., the slice highlighted in yellow in Figure 7a). A circle-finding algorithm was applied to the implant cross-sections to fit a circle and determine the implant center coordinates. Using the center positions obtained at both sides of the implant, a linear interpolation was performed to estimate the implant center for each slice along the implant axis.

For each lacuna, the minimum radial distance between its center of mass and the implant center was then calculated. Because the samples were cylindrical (radius 1.5 mm) and contained an implant of 0.8 mm radius, lacunae located at distances smaller than 0.7 mm or larger than 1.6 mm were excluded from the analysis. For the distance-to-implant analysis, lacunae were grouped into ten radial distance bins, and the median value of each parameter of interest was calculated for each bin.

## Author contributions

**Irene Rodriguez-Fernandez**: Conceptualization, Data curation, Formal analysis, Investigation, Software, Visualization, Writing – original draft. **Thomas Bretschneider**: Conceptualization, Data curation, Formal analysis, Investigation, Software, Visualization, Writing – review & editing. **Goran Lovric**: Investigation, Formal analysis, Software, Writing – review & editing. **Marco Stampanoni**: Supervision, Writing – review & editing. **Omer Suljevic**: Conceptualization, Resources, Writing – review & editing. **Nicole G. Sommer**: Conceptualization, Resources, Writing – review & editing. **Annelie-M. Weinberg**: Conceptualization, Funding acquisition, Resources, Writing – review & editing. **Uwe Y. Schwarze**: Resources, Writing – review & editing. **Lukas Pircher**: Conceptualization, Software, Writing – review & editing. **Christian Hellmich**: Conceptualization, Funding acquisition, Writing – review & editing. **Tilman A. Grünewald**: Conceptualization, Supervision, Writing – review & editing. **Helga C. Lichtenegger**: Conceptualization, Data curation, Funding acquisition, Project administration, Supervision, Writing – review & editing. **Andreas Menzel**: Conceptualization, Data curation, Investigation, Supervision, Writing – review & editing.

## Acknowledgments

We acknowledge the Paul Scherrer Institute PSI for granting experimental time at the X02DA beamline of the Swiss Light Source. We thank Arthur Baroni and Mads Allerup Carlsen for their support during beamtime and Oliver Bunk for fruitful discussions. Views and opinions expressed are those of the author(s) only and do not necessarily reflect those of the European Union or the European Research Council. Neither the European Union nor the granting authority can be held responsible for them.

## Data availability statement

The data that support the findings of this study will be openly available in the PSI Data Catalogue (SciCat [63]) upon publication.

