## Supplementary Information for "The Effects of Physical Exercise on Osteocyte Lacunar Morphology and Orientation around Bioresorbable Implants"

Table S1: Lacunae morphological parameters in newly formed bone. The values represent the mean of all lacunae within one sample with their standard deviation.

| | Lc.N | Lc.N/Lc.T (%) | Stretch | Lc.L1 µm | Lc.L2 µm | Lc.L3 µm | Ratio (L1:L2:L3) | Lc.V $\mu m^3$ | Lc.D $mm^{-3}$ |
|---|---|---|---|---|---|---|---|---|---|
| **2w-nEx** | 120 466 | 24.6 | $0.66 \pm 0.14$ | $16.64 \pm 5.81$ | $9.08 \pm 2.96$ | $5.19 \pm 1.73$ | 3.2:1.75:1 | $470.4 \pm 353.8$ | 60 233 |
| **2w-Ex** | 183 595 | 31.9 | $0.64 \pm 0.14$ | $16.58 \pm 5.08$ | $9.40 \pm 2.61$ | $5.61 \pm 1.61$ | 2.96:1.67:1 | $501.1 \pm 327.6$ | 67 998 |
| **4w-nEx** | 363 207 | 49.2 | $0.64 \pm 0.14$ | $15.59 \pm 4.40$ | $8.90 \pm 2.15$ | $5.06 \pm 1.34$ | 3.2:1.75:1 | $388.0 \pm 240.3$ | 73 227 |
| **4w-Ex** | 263 878 | 36.8 | $0.64 \pm 0.14$ | $15.44 \pm 4.76$ | $8.79 \pm 2.15$ | $5.19 \pm 1.35$ | 2.96:1.67:1 | $392.2 \pm 255.3$ | 74 123 |
| **6w-nEx** | 326 108 | 38.2 | $0.68 \pm 0.13$ | $15.44 \pm 4.61$ | $8.38 \pm 2.22$ | $4.42 \pm 1.29$ | 3.49:1.9:1 | $321.1 \pm 228.7$ | 69 385 |
| **6w-Ex** | 193 450 | 29.9 | $0.65 \pm 0.13$ | $15.13 \pm 4.34$ | $8.52 \pm 2.08$ | $4.91 \pm 1.23$ | 3.08:1.74:1 | $347.3 \pm 213.5$ | 71 384 |

Lc.N - number of lacunae
Lc.N/Lc.T - percentage of lacunae of the total lacunae in the full sample
Lc.L1 - long axis µm
Lc.L2 - medium axis µm
Lc.L3 - short axis µm
Lc.V - lacunar volume $\mu m^3$
Lc.D - lacunar density $mm^{-3}$

Table S2: Lacunae morphological parameters in previously existing bone. The values represent the mean and standard deviation. (a) Closer than 200 µm from the implant. (b) Farther than 200 µm from the implant. (c) Total.

**(a) Closer than** 200 µm

| | Lc.N | Stretch | Lc.L1 µm | Lc.L2 µm | Lc.L3 µm | Ratio (L1:L2:L3) | Lc.V $\mu m^3$ | Lc.D $mm^{-3}$ |
|---|---|---|---|---|---|---|---|---|
| **2w-nEx** | 70 468 | $0.69 \pm 0.14$ | $14.85 \pm 4.87$ | $7.67 \pm 2.36$ | $4.38 \pm 1.23$ | 3.39:1.75:1 | $282.4 \pm 218.5$ | 43 499 |
| **2w-Ex** | 47 965 | $0.67 \pm 0.14$ | $15.54 \pm 5.27$ | $8.36 \pm 2.53$ | $4.88 \pm 1.42$ | 3.18:1.71:1 | $367.5 \pm 277.9$ | 58 494 |
| **4w-nEx** | 124 337 | $0.65 \pm 0.14$ | $15.50 \pm 5.77$ | $8.74 \pm 2.50$ | $4.83 \pm 1.51$ | 3.21:1.81:1 | $371.3 \pm 262.4$ | 56 775 |
| **4w-Ex** | 145 586 | $0.65 \pm 0.14$ | $15.29 \pm 5.44$ | $8.66 \pm 2.53$ | $4.70 \pm 1.51$ | 3.25:1.84:1 | $358.1 \pm 270.1$ | 55 146 |
| **6w-nEx** | 94 441 | $0.69 \pm 0.14$ | $15.38 \pm 5.58$ | $7.92 \pm 2.50$ | $4.33 \pm 1.29$ | 3.55:1.83:1 | $299.6 \pm 231.7$ | 59 026 |
| **6w-Ex** | 115 179 | $0.65 \pm 0.14$ | $15.29 \pm 5.05$ | $8.55 \pm 2.45$ | $4.83 \pm 1.43$ | 3.17:1.77:1 | $360.0 \pm 258.5$ | 63 988 |

**(b) Farther than** 200 µm

| | Lc.N | Stretch | Lc.L1 µm | Lc.L2 µm | Lc.L3 µm | Ratio (L1:L2:L3) | Lc.V $\mu m^3$ | Lc.D $mm^{-3}$ |
|---|---|---|---|---|---|---|---|---|
| **2w-nEx** | 298 939 | $0.68 \pm 0.13$ | $14.97 \pm 4.11$ | $7.93 \pm 2.11$ | $4.62 \pm 1.13$ | 3.24:1.72:1 | $305.1 \pm 203.5$ | 72 912 |
| **2w-Ex** | 343 892 | $0.67 \pm 0.13$ | $15.21 \pm 4.30$ | $8.26 \pm 2.10$ | $4.83 \pm 1.19$ | 3.15:1.71:1 | $338.7 \pm 220.8$ | 63 449 |
| **4w-nEx** | 250 831 | $0.66 \pm 0.13$ | $15.40 \pm 4.16$ | $8.66 \pm 2.13$ | $4.88 \pm 1.19$ | 3.16:1.77:1 | $358.6 \pm 211.8$ | 61 781 |
| **4w-Ex** | 307 937 | $0.65 \pm 0.13$ | $14.58 \pm 4.48$ | $8.38 \pm 2.18$ | $4.54 \pm 1.26$ | 3.21:1.85:1 | $315.3 \pm 223.8$ | 61 711 |
| **6w-nEx** | 432 495 | $0.70 \pm 0.13$ | $14.90 \pm 4.27$ | $7.59 \pm 1.90$ | $4.33 \pm 1.02$ | 3.44:1.75:1 | $266.3 \pm 165.5$ | 71 724 |
| **6w-Ex** | 339 397 | $0.67 \pm 0.13$ | $14.98 \pm 4.46$ | $8.18 \pm 2.07$ | $4.60 \pm 1.15$ | 3.26:1.78:1 | $312.5 \pm 201.1$ | 62 047 |

**(c) Total**

| | Lc.N | Lc.N/Lc.T (%) | Stretch | Lc.L1 µm | Lc.L2 µm | Lc.L3 µm | Ratio (L1:L2:L3) | Lc.V $\mu m^3$ | Lc.D $mm^{-3}$ |
|---|---|---|---|---|---|---|---|---|---|
| **2w-nEx** | 369 104 | 75.4 | $0.68 \pm 0.13$ | $14.95 \pm 4.26$ | $7.88 \pm 2.16$ | $4.57 \pm 1.15$ | 3.27:1.72:1 | $300.8 \pm 206.6$ | 64 529 |
| **2w-Ex** | 392 283 | 68.1 | $0.66 \pm 0.13$ | $15.25 \pm 4.47$ | $8.27 \pm 2.16$ | $4.84 \pm 1.22$ | 3.15:1.71:1 | $342.0 \pm 228.8$ | 62 866 |
| **4w-nEx** | 375 320 | 50.8 | $0.66 \pm 0.13$ | $15.44 \pm 4.75$ | $8.68 \pm 2.26$ | $4.86 \pm 1.30$ | 3.18:1.79:1 | $362.6 \pm 230.0$ | 60 051 |
| **4w-Ex** | 453 192 | 63.2 | $0.65 \pm 0.13$ | $14.81 \pm 4.82$ | $8.47 \pm 2.30$ | $4.59 \pm 1.35$ | 3.23:1.85:1 | $329.0 \pm 240.5$ | 59 396 |
| **6w-nEx** | 526 585 | 61.8 | $0.70 \pm 0.13$ | $14.99 \pm 4.54$ | $7.65 \pm 2.02$ | $4.33 \pm 1.07$ | 3.46:1.77:1 | $272.2 \pm 179.6$ | 69 015 |
| **6w-Ex** | 454 522 | 70.1 | $0.66 \pm 0.13$ | $15.06 \pm 4.63$ | $8.27 \pm 2.18$ | $4.66 \pm 1.23$ | 3.23:1.77:1 | $324.5 \pm 218.1$ | 62 349 |

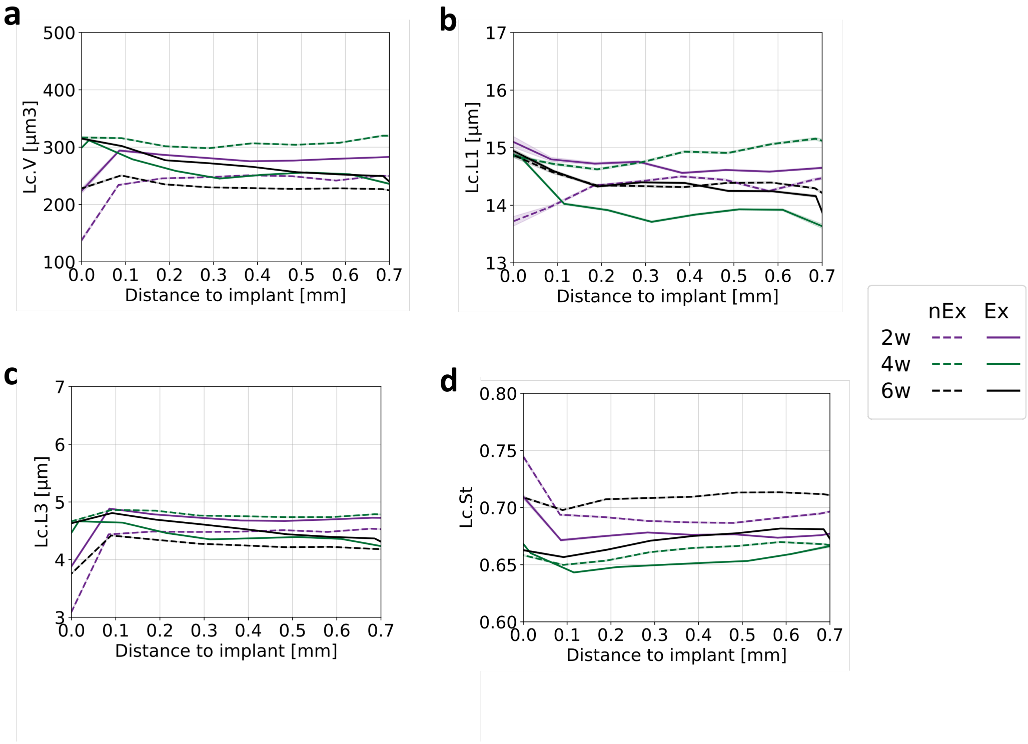


Figure S1: Lacunar morphological properties as a function of distance from the bone–implant interface for previously existing bone. (a) Lacuna volume (Lc.V), (b) long axis (Lc.L1), (c) short axis (Lc.L3), and (d) stretch.

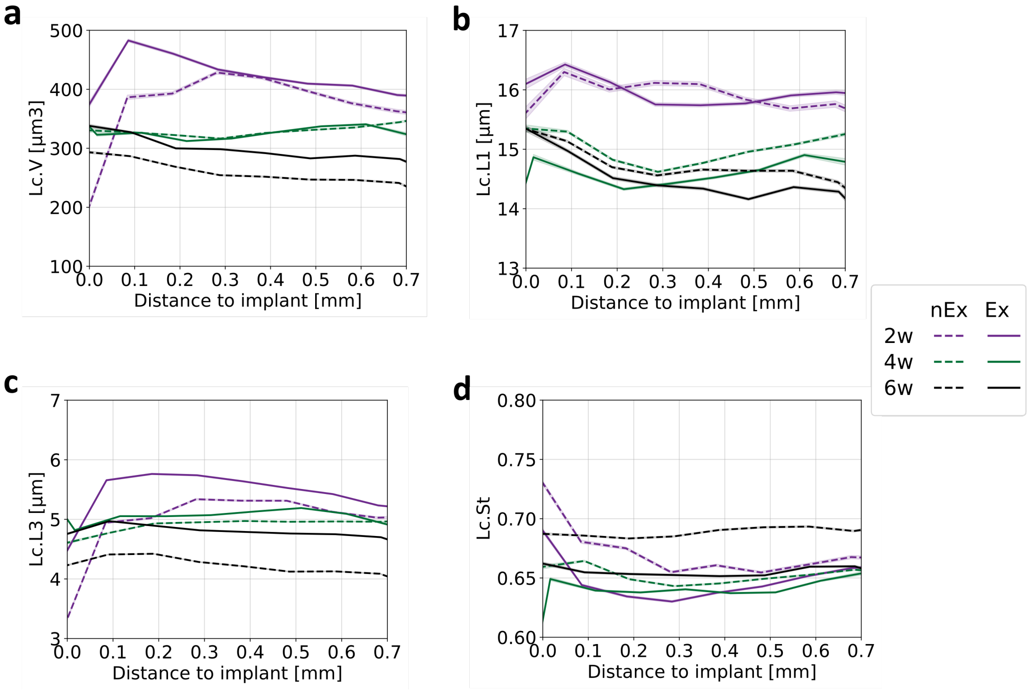


Figure S2: Lacunar morphological properties as a function of distance from the bone–implant interface for newly formed bone. (a) Lacuna volume (Lc.V), (b) long axis (Lc.L1), (c) short axis (Lc.L3), and (d) stretch.

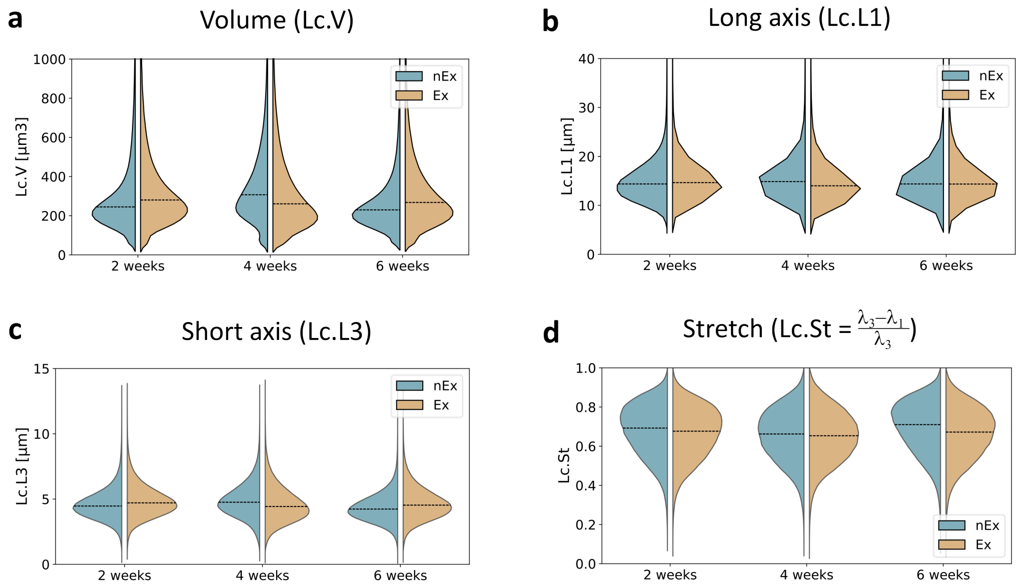


Figure S3: **Distributions of lacunar morphological properties in previously existing bone.** Plots comparing the distributions of lacunar volume (a), Lc.L1 (long axis) (b), Lc.L3 (short axis) (c), and stretch (d). The dashed line represents the median value of the distribution.
